# Spiders as surrogates of biodiversity in a geothermal area of northern Puebla, México

**DOI:** 10.1101/2022.08.23.505016

**Authors:** Luis Carlos Hernández-Salgado, Guillermo Romero, Jesus Rodriguez-Canseco, Krystal Gonzalez-Estupiñan

## Abstract

The word biodiversity has gained popularity for more than 30 years; and we used this term for all aspects of biological diversity, from the richness of taxa, genetic variation, and ecosystem complexity. Thoroughly measuring biodiversity would be impractical due to many existing species and the effort that their identification would represent. Therefore, we need to find suitable surrogates or substitute taxa, which we can monitor because they represent broader biodiversity, helping us in ecological monitoring or conservation program planning. We tested the richness of spiders to measure their ability to estimate total spider richness at a site with geothermal influence. We use data from six locations with geothermal effect in the northern highlands of Puebla, Mexico. The genera richness of spiders was a good substitute for the total species richness, and the opposite happened with the family richness. Therefore, we recommend using this taxonomic level as a substitute to predict spider species richness or to evaluate and classify sites according to their importance for conservation. Here, we present the first spiders biodiversity data of a geothermal area and the use of spiders as surrogates in sites with geothermal condition.

## Introduction

The term biodiversity has gained popularity for more than 30 years. We use this term for all aspects of biological diversity, including taxa richness, genetic variation, and ecosystem complexity (Magurran, 2004). Due to many existing species and the effort that their identification represents, the measurement of total biodiversity is unreasonable. Therefore, for effective ecological monitoring, the estimations of biodiversity or support in conservation programs are necessary to find surrogates or substitute taxa that can indicate the state of the biodiversity or environment (Anderson et al., 2011; Moreno & Sanchez-Rojas, 2007).

Biological indicator species are species or groups of species that indicate the environmental state. They help in the early warning of environmental changes and monitor ecosystem stressors or identify site diversity (Kovács, 1992; McGeoch, 1998). Biological indicator species have a narrow amplitude for one or more environmental factors. When present, it indicates a particular environmental condition or set of conditions, for example, habitat fragmentation or pollution (Allaby, 2014).

The overwhelming pressure imposed by human activities, such as energy production, has impact on natural systems. This impact mainly depends on the technology used by the output (Rybach, 2003). The identification of the effects that such activity has on the environment is essential to evaluate the environmental impact.

The development of geothermal energy causes gas emission and effluent water, which produces chemical pollutants discharges into the environment (Kristmannsdóttir & Ármannsson, 2003; Rybach, 2003; Thórhallsdóttir, 2007), with substantial changes in the biological function of the ecosystem in which they are poured (Azam et al., 2015).

Physical and chemical analyses provide data on the presence, levels, and degradation of environmental to evaluate the level of environmental pollution produced by the development of this type of energy. However, these parameters do not reflect the environmental stress that this pollution or degradation causes in living organisms or their effects due to this stress (Omar, 2010). We need to search for economic and easy-to-apply methods that allow the early detection of environmental disturbances that could endanger biodiversity and human health. One option to address these issues is surrogates’ use (Gonzalez Zuarth & Vallarino, 2014).

Microalgae and cyanobacteria (Nuñez Zarco, 2016), birds (Dmowski, 1999; Frederick, Spalding, & Dusek, 2002), and bats (Zukal, Pikula, & Bandouchova, 2015) already been used as surrogates. Terrestrial and aquatic arthropods, constitute a substantial proportion of the species richness and biomass in the ecosystems, also play an essential role in their functioning, nevertheless has been rarely used as surrogates (Hodkinson & Jackson, 2005; Mauricio da Rocha, De Almeida, Lins, & Durval, 2010; McGeoch, 1998). Within this group, spiders are an excellent option to use as environmental, ecological, and biodiversity indicators because they are susceptible to environmental disturbances, are highly diverse, abundant, and have a wide variety of ecological niches, and are mainly predators of the insect group (Gerlach, Samways, & Pryke, 2013).

We hypothesized that the higher spider taxa will provide us with information on the total diversity of spiders, and to test our hypothesis, we will evaluate the feasibility of the higher spider taxa as indicators of biodiversity (also called “biodiversity substitutes”) in a geothermal zone in Puebla, México. The contribution of this study will allow us to use spiders as a fast, effective, and low-cost tool for ecological studies in geothermal sites, thereby supporting the realization of a management proposal and assessing priority areas for the conservation of these organisms.

## Methods

### Study site

The geothermal zone of Acoculco (589421.57 E, 2203181.12 N) borders the state of Hidalgo and is in the town of Chignahuapan in the state of Puebla 85 km northwest of the city of Puebla and 65 km southwest of the city of Pachuca, Hidalgo, México. The elevation of the geothermal zone is between 2,800 and 2,900 meters above sea level. (Hiriart Le Bert, 2011). The Acoculco area covers an area of 1,290 km ^2^ and includes 39 springs and some CO_2_ emissions. It is in the border zone between the Mexican Volcanic Belt and the Madre Oriental mountain range, where tertiary and quaternary volcanic rocks emerge. A pine forest surrounds the geothermal site of Acoculco, and the emanations are in a grassland (Fig. 1) (Quinto, Santoyo, Torres, Gonzalez, & Castillo, 1995).

**Figure. 1.**
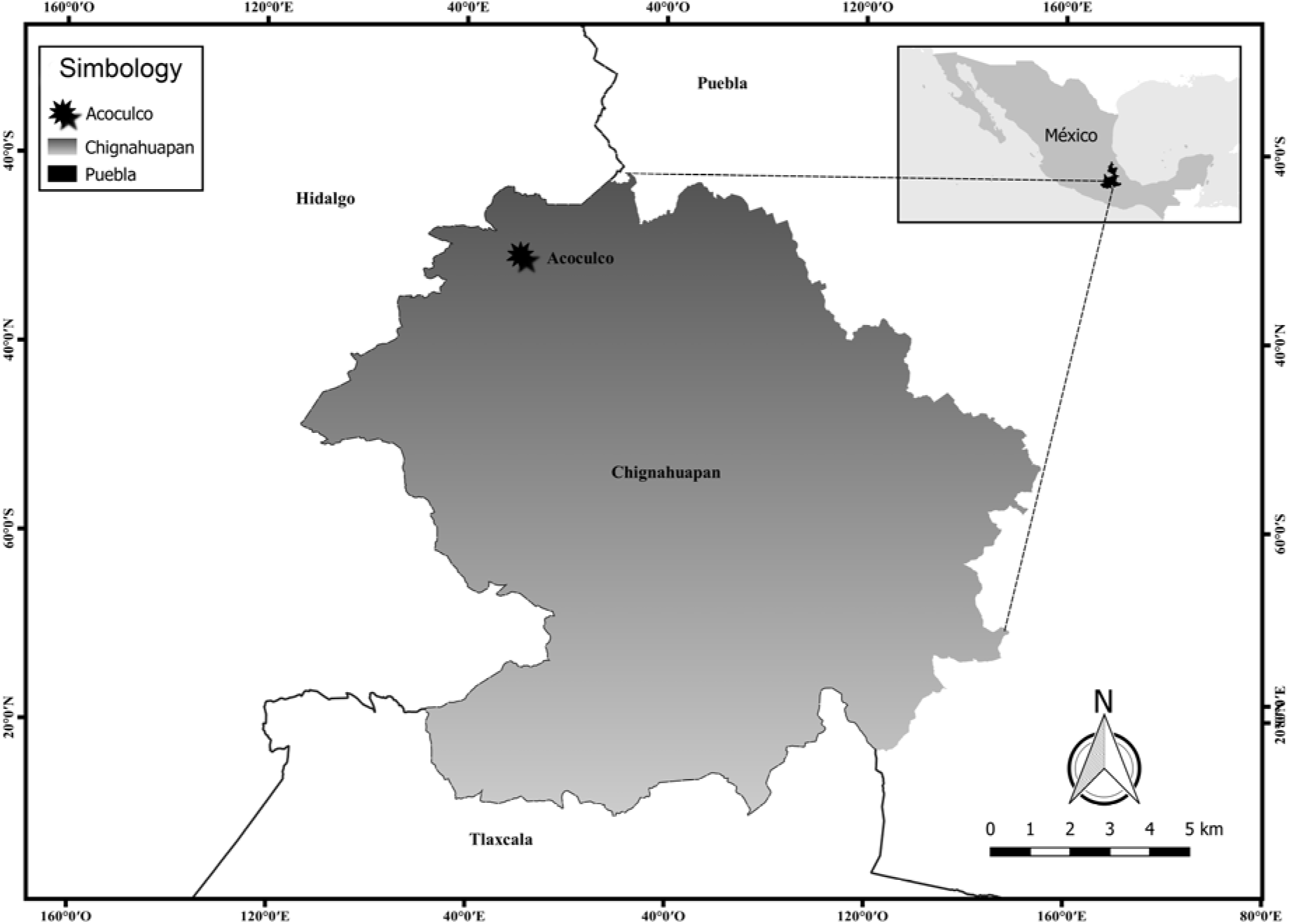
Map of the geothermal zone of Acoculco, Municipality of Chignahuapan, Puebla México.

### Fieldwork

The sampling was carried out during the rainy season from November 19, 2018, to November 30, 2018, in the geothermal area of Acoculco. Sampling lasted for a total of 12 days. We carried out sampling in six locations in the geothermal zone; we also considered vegetation types in each location. The six sites sampled were: Cruz Colorada (CCA) (forest and grassland), Cuautelolulco (CTL) (forest and grassland), La Campana (CLA) (forest and grassland), Tenancingo (TEN) (scrub), Jonuco (JON) (forest) and Michac (MIC) (oak forest) (Fig. 2). Geothermal emanations are in the town of Cruz Colorada (Fig. 3). In each location, we established two 50-meter transects (12 in total). We selected five sampling points of five meters in diameter in each transect, each separated by 10 meters. We made spider collects at each point with manual searches in the soil and vegetation and with the help of a blanket placed under the vegetation foliage, which we struck with a stick to make fall to the organisms captured with the help of brushes and entomological tweezers. Two collectors did the capture of spiders. The specimens were placed in 70% alcohol to transport them to the laboratory (Marquez Luna, 2005) and identified to family, genera, and species level when possible. We deposited the collected spiders in the arthropod collection of the Science Faculty of the Autonomous University of Baja California. We considered each sampling transect as a different sample; we obtain a total of 12 samples.

**Figure. 2.**
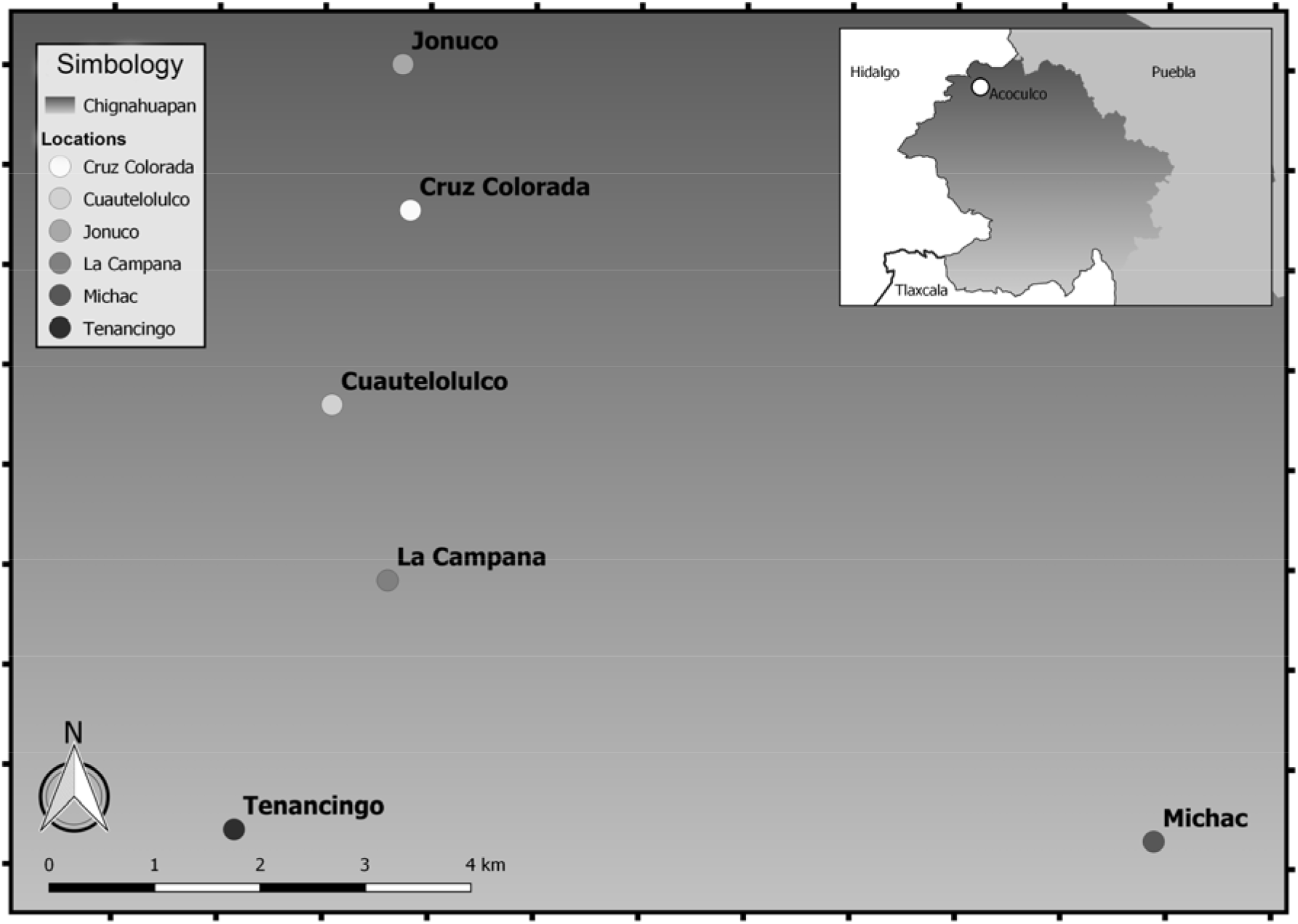
Map with the sampled locations in the geothermal area of Acoculco, Puebla, México.

**Figure. 3.**
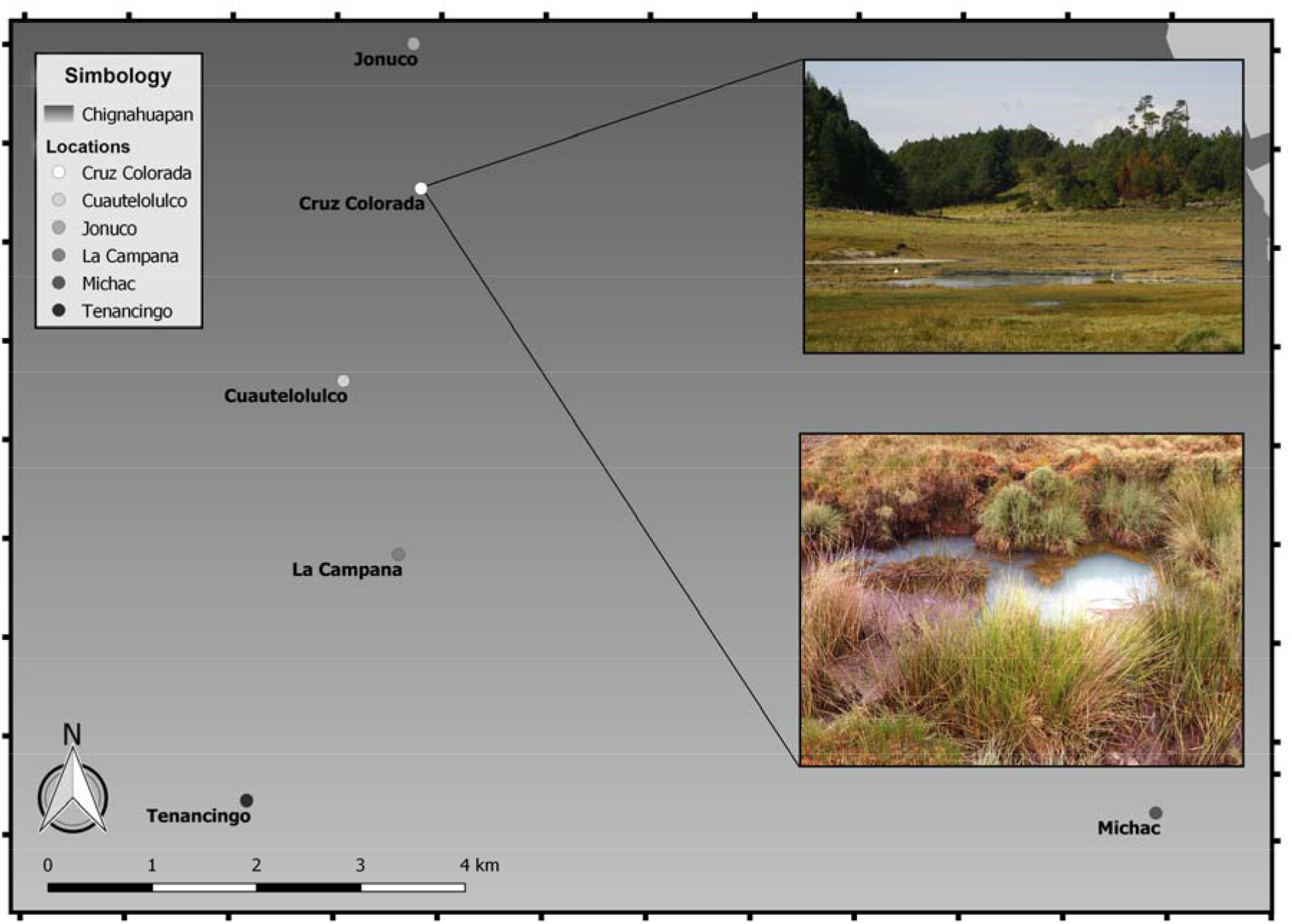
Map illustrating the geothermal emanations present in the town of Cruz Colorada.

### Statistical procedures

To evaluate the integrity and statistical significance of the sampling, we obtained the total richness with the help of the EstimateS version 9.0 program (Colwell, 2013); using two non-parametric richness estimators Chao 1 and ACE (Magurran, 2004), and with the data obtained, we construct species accumulation curves for the entire sampling data. The proportion of “Singletons” and “Doubletons” was calculated as an additional tool to evaluate the sampling (Coddington, Griswold, Silva Davila, Efrain, & Larcher, 1991). We calculate two indexes with the software Past (Hammer et al.,2001), The Shannon-Wiener (H’) and Simpson (D), because they have different sensitivity, the Shannon-Wiener index is sensitive to rare species. In contrast, the Simpson index is sensitive to changes in the abundance of the most common species.

We perform regression analyses to test if higher spider taxa richness can function as substitutes for diversity. The higher taxa analyzed were families and genera; these were taken as independent variables, while the species were dependent variables. Linear, exponential, and logarithmic regressions were tested, and we chose those with the highest regression coefficient. We used the percentage of the variance explained by the independent variable (r^2^) and the scatter plots’ visual evaluation as an adjustment measure, subrogation reliability, and predictive power (Cardoso, Silva, de Oliveira, & Serrano, 2004).

According to the methodology used by Cardoso (2004), we tested two approaches for prioritization and ranking of sites for conservation. A scoring approach uses the raw number of taxa represented in each site as the sole value for ranking. We used the Spearman rank correlation index to test for surrogacy reliability in the scoring of sites. We use scatter plots of family and genus richness versus species richness ranking sites to inspect reliability visually. We furthermore tested a more efficient iterative approach of conservation priority ranking. For each of the considered taxonomic levels (family, genus, or species), we first chose the richest site, and from it, in a stepwise manner, the one site that would further raise the number of represented taxa was added to the set of sites to be considered for protection. In the case of ties, we chose the richest site in the respective taxa; the objective is to check what species’ proportion can be protected by using the same number of sites that protect all considered higher taxa. By doing so, we could test the effect of using higher taxa for choosing a near-minimum set of sites that potentially preserves the maximum number of species.

According to the methodology used by Cardoso (2004b), we perform regression analyses to test if we can use a single family or a group of spider families to predict all species richness. The percentage of the variance explained by the independent variable (*r^2^*), and we use the scatter charts’ visual evaluation as a measure of fit, surrogacy reliability, and predictive power. We select the family with the highest regression coefficient, and we took it as a starting point to later add the families that manage to increase the value of the regression coefficient; we carried out this procedure until the smallest group of possible families was reunited. This procedure was carried out with the data of all the families found. As with the higher taxa, we tested two approaches to determine the priority and ranking of sites for conservation. We use the Spearman correlation index and taxa accumulation curves by the site to verify that proportion of the total spider species can be protected using the same number of sites that protect the species of taxa (families or genera) previously analyzed.

## Results

### Diversity analysis and Sampling efficiency

We collected 317 spiders belonging to 11 families, 27 genera, and 49 species or morphospecies. We identified all spiders at least up to genus level, that was necessary for all analyzes. Of the 11 families recorded in the entire sample, the most abundant were Theriididae with 64 individuals (20%), Lycosidae and Tetragnathidae with 56 individuals (18%), and Anyphaenidae and Salticidae with 33 individuals (10%); in contrast, the least abundant families were Agelenidae and Thomisidae, with 11 individuals (2%), and Zoropsidae with only three individuals (1%). The most abundant species in the entire sample were *Pardosa sternalis* (Thorell, 1877) with 44 individuals (14%), *Anhypaena catalina* (Platnick, 1974), and *Tetragnatha elongata* (Walckenaer, 1841) with 33 individuals (10%), *Theridion trepidum* (O. Pickard-Cambridge, 1898) with 26 individuals, and *Latrodectus mactans* (Fabricius, 1775) with 18 individuals (6%).

The families with the highest species richness were Araneidae with 11 species, Salticidae and Lycosidae with eight species, and Theriididae with five species. The most abundant species in the Cruz Colorada locality were *P. sternalis* and *Dictyna puebla* (Gertsch & Davis, 1937) (13%), *Tetragnathidae sp1* (12%), and *T. elongata* and *A. catalina* (10%). In Cuautelolulco the most abundant species were *P. sternalis* (18%), *L. mactans* (15%), *Salticidae sp1* (13%) and *A. catalina* (10%). In the town of La Campana, the most abundant species were *Salticidae sp1* and *Paraphidippus aurantius* (Lucas, 1833) (21%), and *T. elongata* (14%). In Jonuco the most abundant species were *T. elongata* (21%), *T. trepidum* (15%), *A. catatalina* (13%) and *Neoscona sp1* (Simon, 1864) (12%). In Michac the most abundant species were *T. trepidum* (24%) and *P. sternalis* (24%), *A. catalina* and *Zorocrates unicolor* (Banks, 1901) (14%) and *Salticidae sp1* (10%). In the town of Tenancingo the most abundant species were *P. sternalis* (19%), *T. trepidum*, and *T. elonogata* (16%). The list of spiders reported for this work can be seen in the table 1.

**Table 1.**
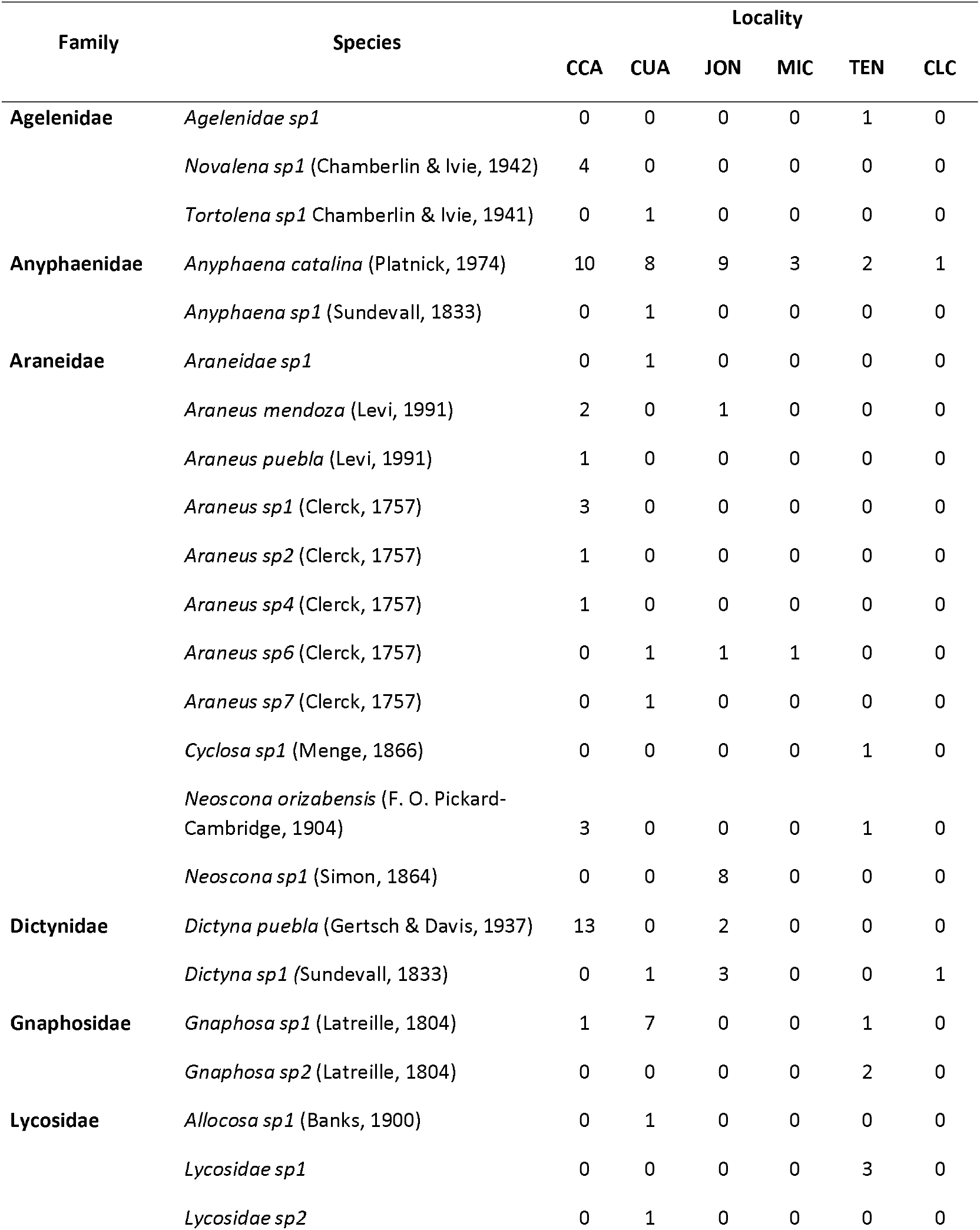
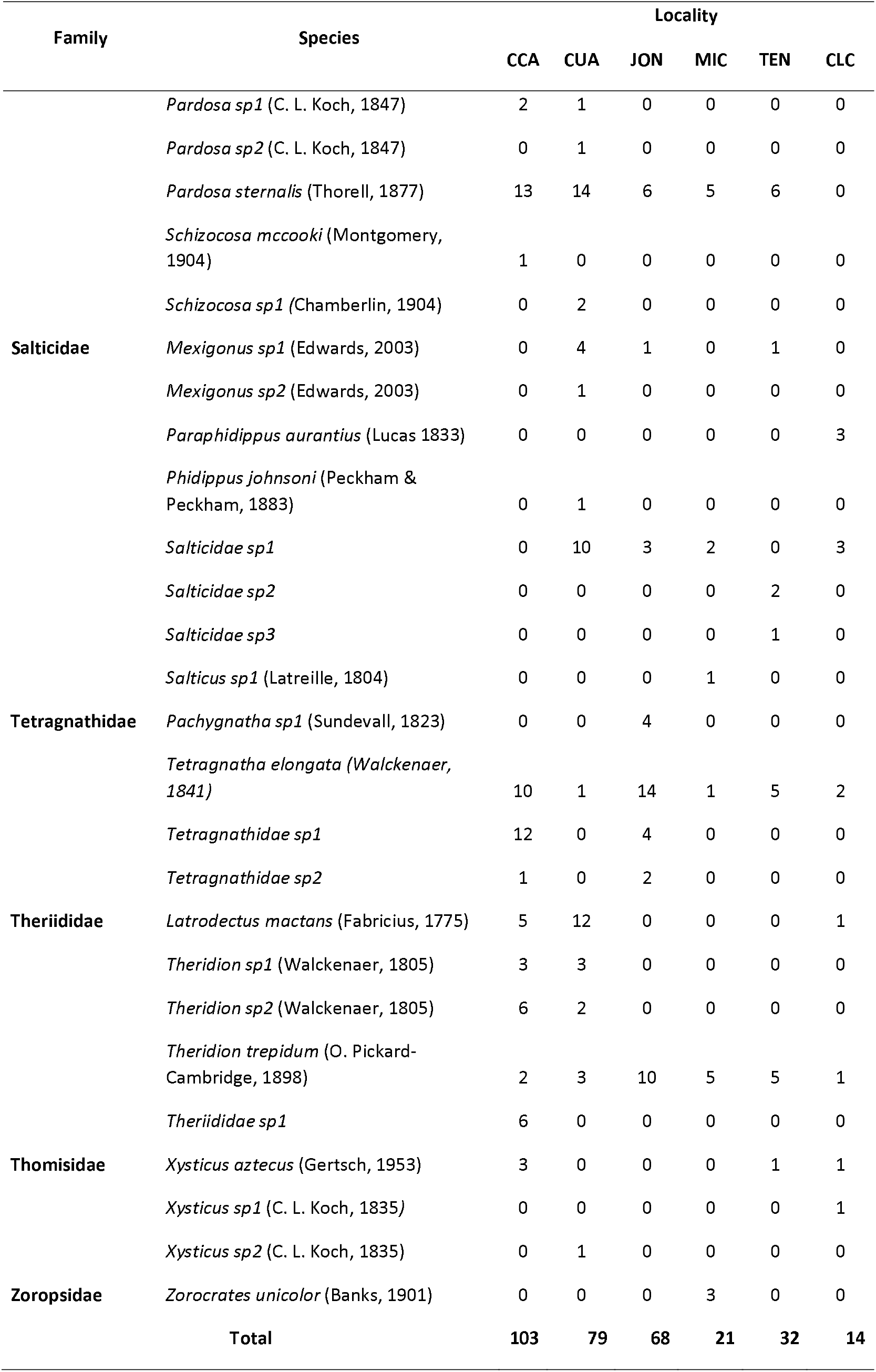
List of the abundance of species and morphospecies of spiders found in the six localities of the geothermal area of Acoculco, Puebla. Localities: Cruz Colorada (CCA), Cuautelolulco (CUA), Jonuco (JON), Tenancingo (TEN), Michac (MIC) and La Campana (CLC).

The Chao1 values indicate a lack in the sample of 41% of the richness, while the ACE estimator suggests a lack of 25%. We obtain an Integrity of 53% with Chao1 and 70% with ACE. According to the Chao1 estimator, 42 species were missing, and according to the ACE estimator, only 20 were missing (Fig. 4). No estimator managed to arrive at the asymptote. We obtain 19 Singletons and 4 Doubletons.

**Figure 4.**
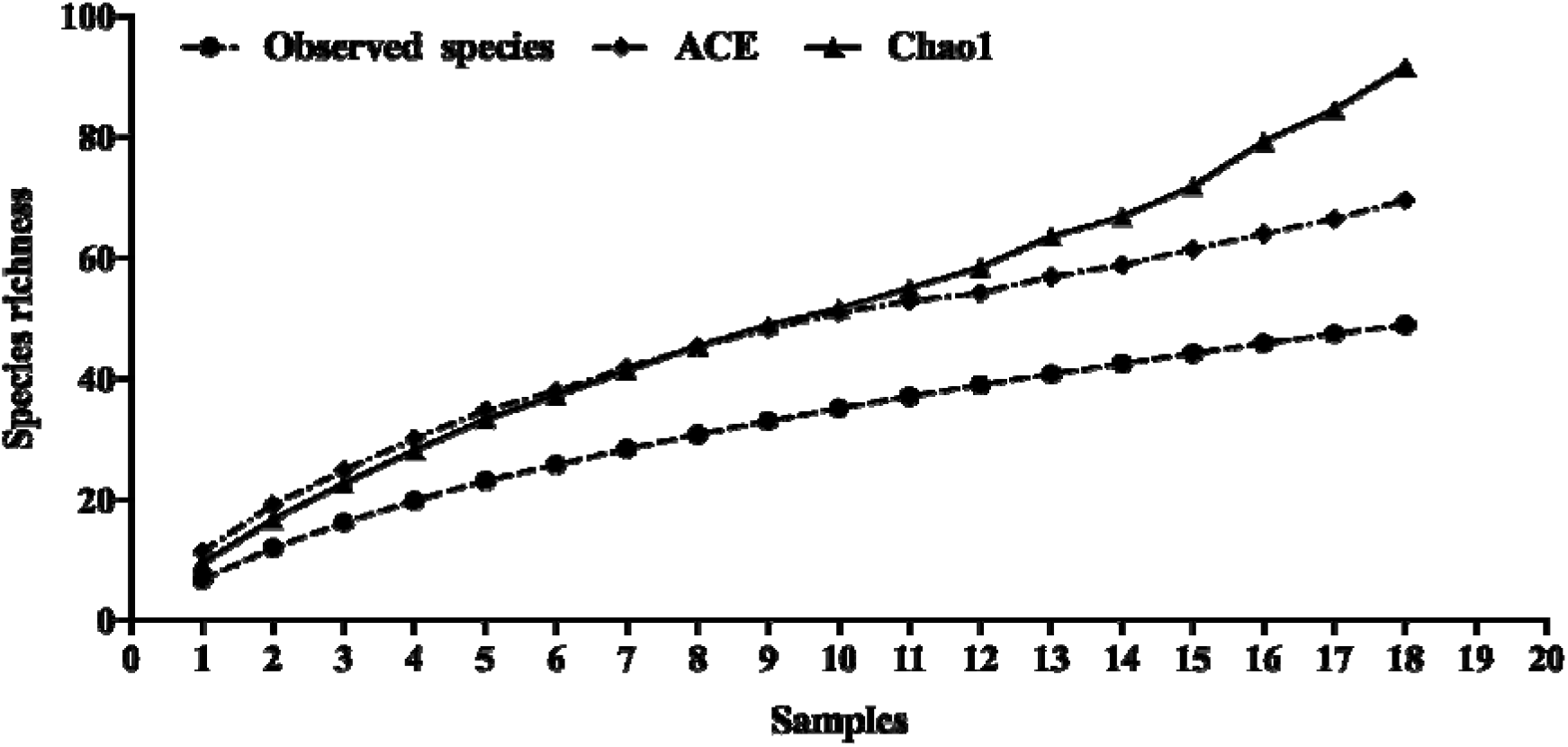
Species accumulation curves for the six sampling locations with the estimators, Chao 1 and ACE.

The Shannon-Wiener index was higher in the locality of Cruz Colorada (H’ = 2,742) and had a similar trend in Cuautelolulco (H’ = 2,661); Tenancingo (H’ = 2,394) and Jonuco (IT = 2,343) also presented a similar trend; La Campana (H’ = 2,069) and Michac (H’ = 1,898) had the lowest values. The Simpson index shows a greater dominance in Cruz Colorada and Cuautelolulco and not so for the remaining localities (Table 2)

**Table 2.**
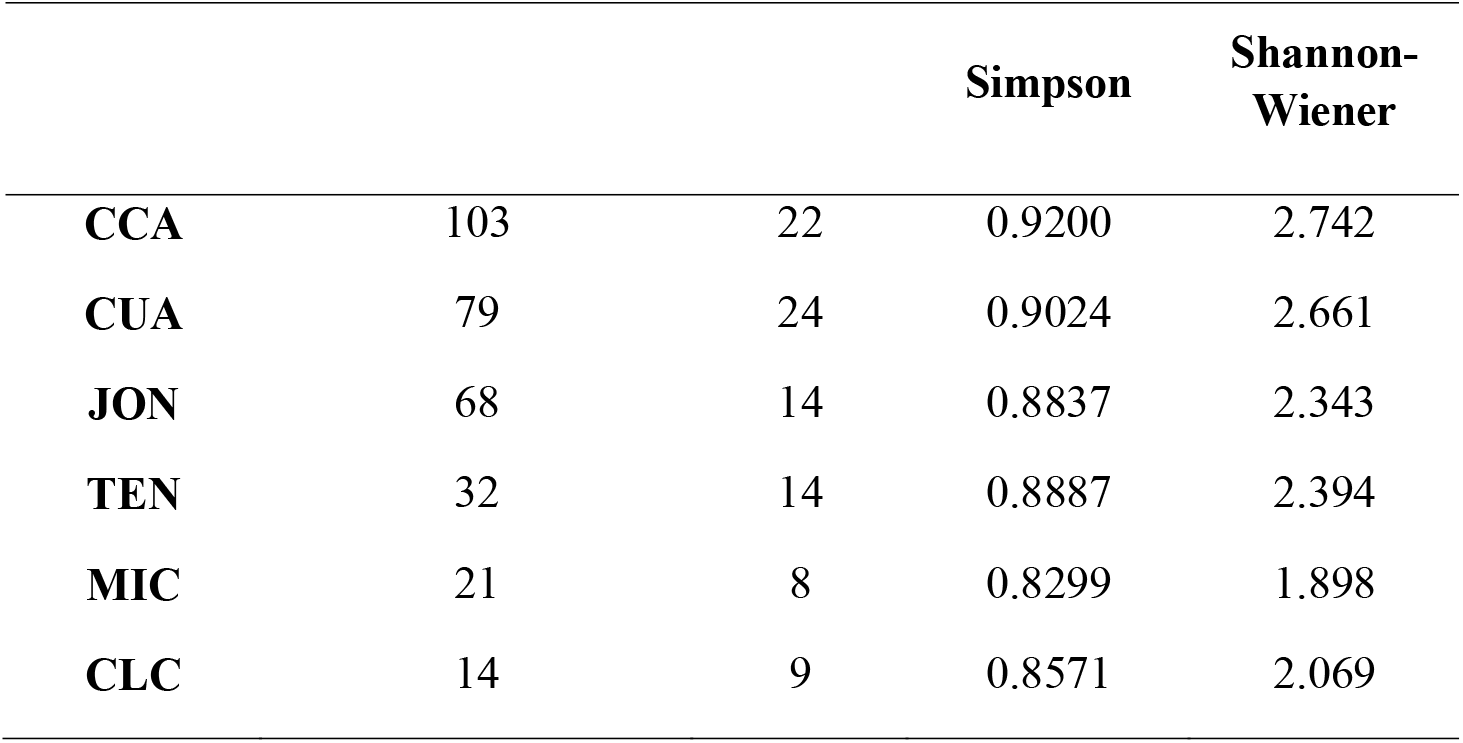
Data on Abundance, richness, Simpson index and Shannon-Wiener, index.

### Higher taxa as surrogates for diversity

We choose the highest regression coefficient value after fitting the regression types to family and genus taxonomic levels. We found a linear relationship for both taxonomic levels. We found a significant correlation between families and species (n = 6, p = 0.0270) with an *r^2^* value of 0.744 (Fig. 5a) and we also found a significant correlation between genus and species (n = 6, p = 0.0013) with a value of *r^2^* of 0.9411 (Fig. 5b). Sites with a similar number of families do not have a similar number of species; the same happens whit the genera. According to the regression analysis, genera’s richness seems to have a greater predictive power since it has a higher value of *r^2^*.

**Figure. 5.**
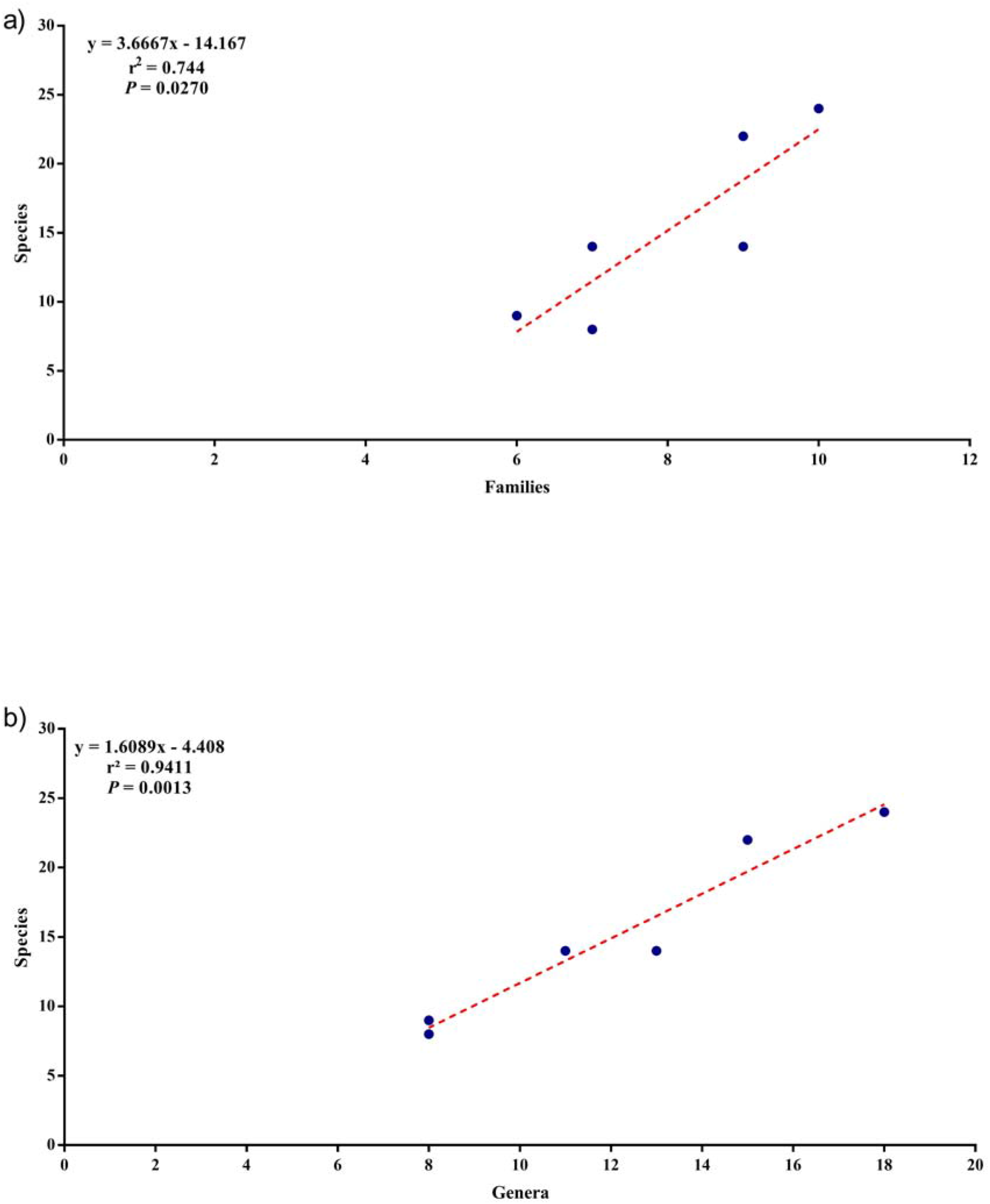
(a) Linear relationship between the richness of families and spider species in the six locations sampled; (b) Linear relationship between the richness of genera and spider species in the six locations sampled.

When classifying sites according to their richness of taxa, we find families to have a low predictive power of species-based site ranking (Table 3). However, they have a high Spearman correlation value of 0.8359. When examining the scatter plot (Fig 6a), we could also observe the spider family surrogacy approach’s low reliability. On the other hand, the spider genus level seems to rank the sites in the same way as species do, with a Spearman value of 0.9706 (Fig. 6b).

**Table 3.**
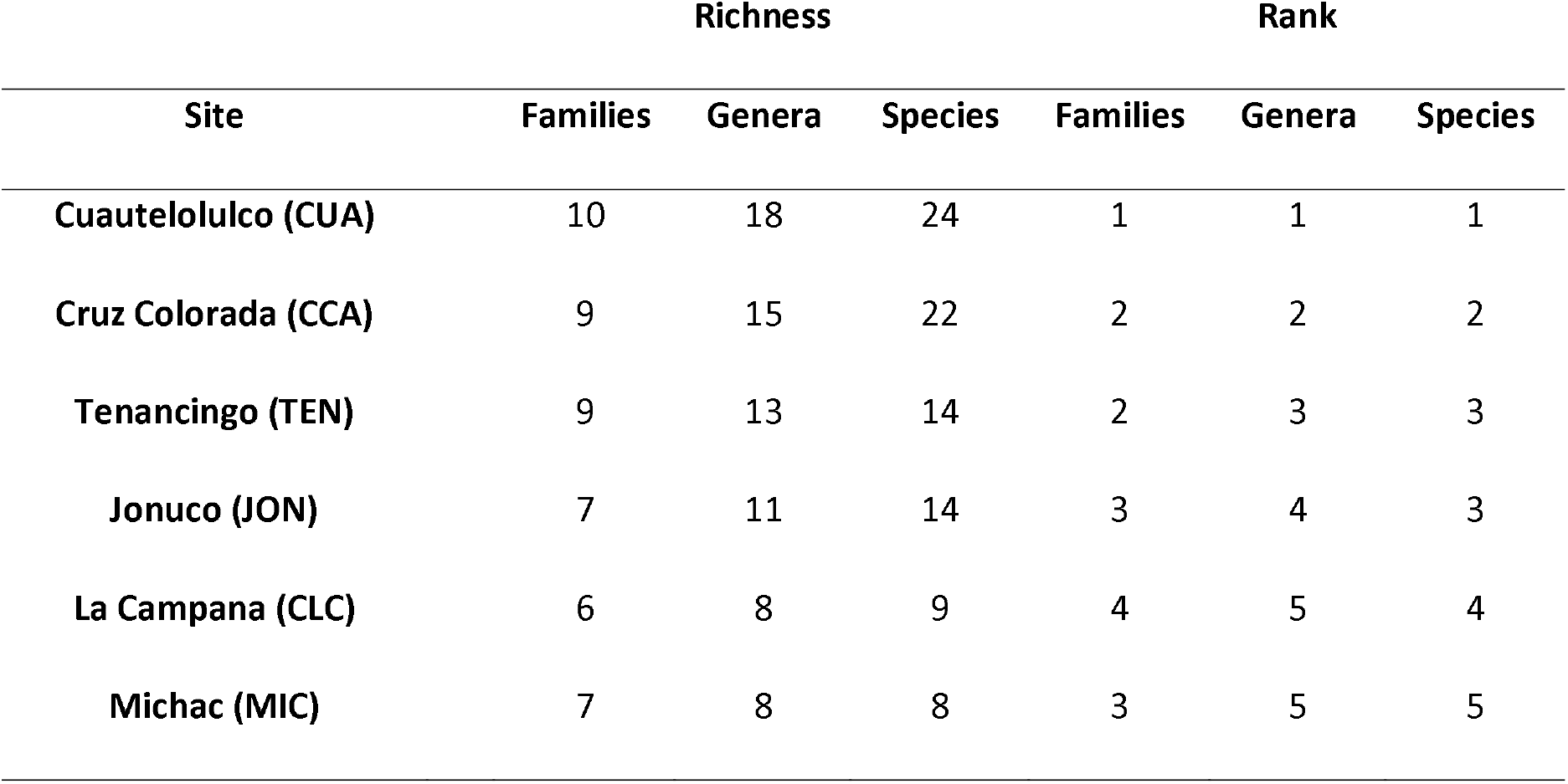
Taxa richness of sampled sites and respective ranking in geothermal sites in Acoculco, Puebla, México.

**Figure 6.**
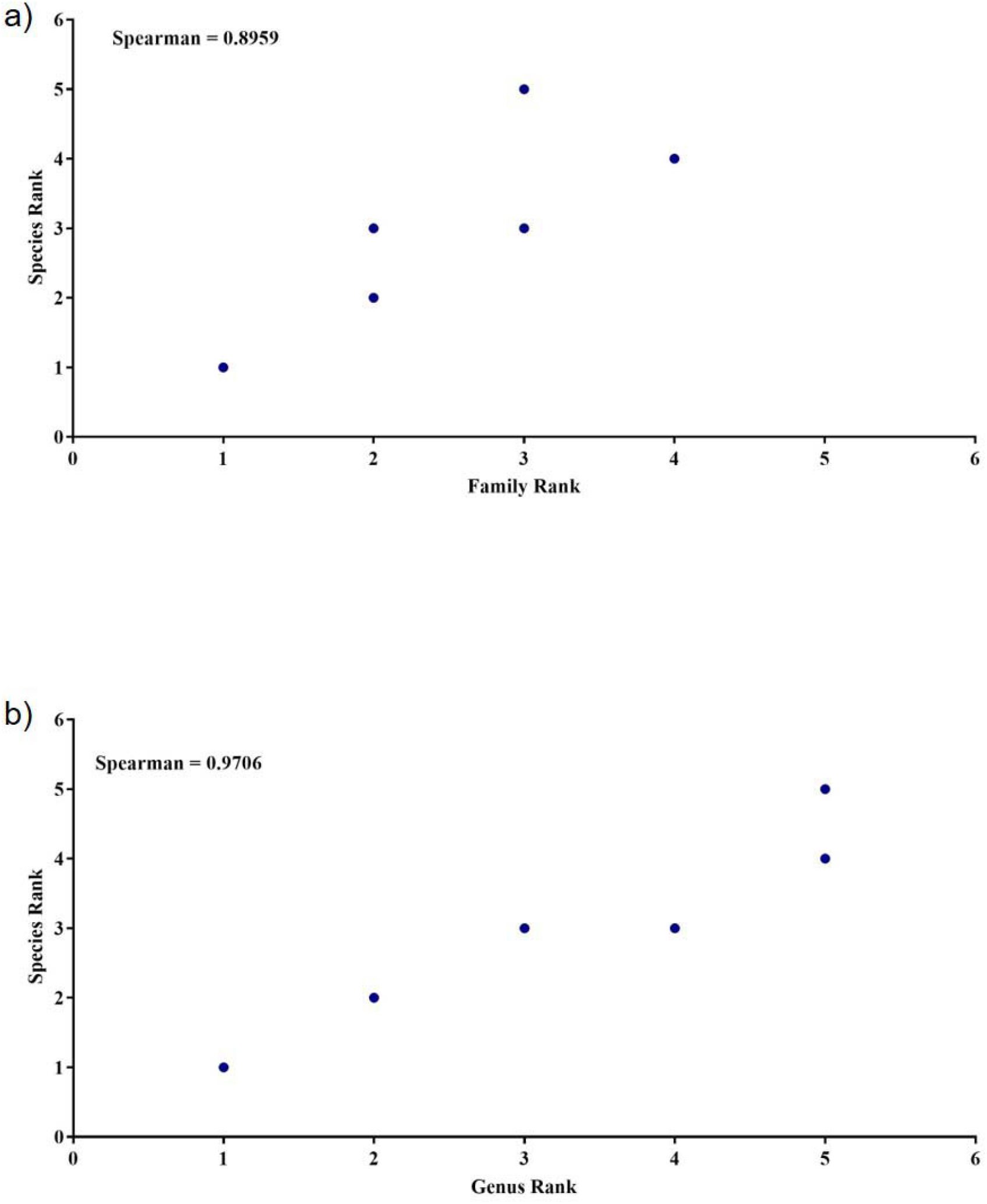
(a) Comparing site ranking according to family and species richness; (b) comparison of site ranking according to genus and species richness.

According to the taxa accumulation curves, five sites (83%) are sufficient to include all species and all genera, and only four sites (67%) for all families. The number of sites needed to have all families is sufficient to protect 90% of the species, whereas, when using the number of sites necessary for all genera, it can be sufficient to protect 100% of the species (Fig. 7).

**Figure 7.**
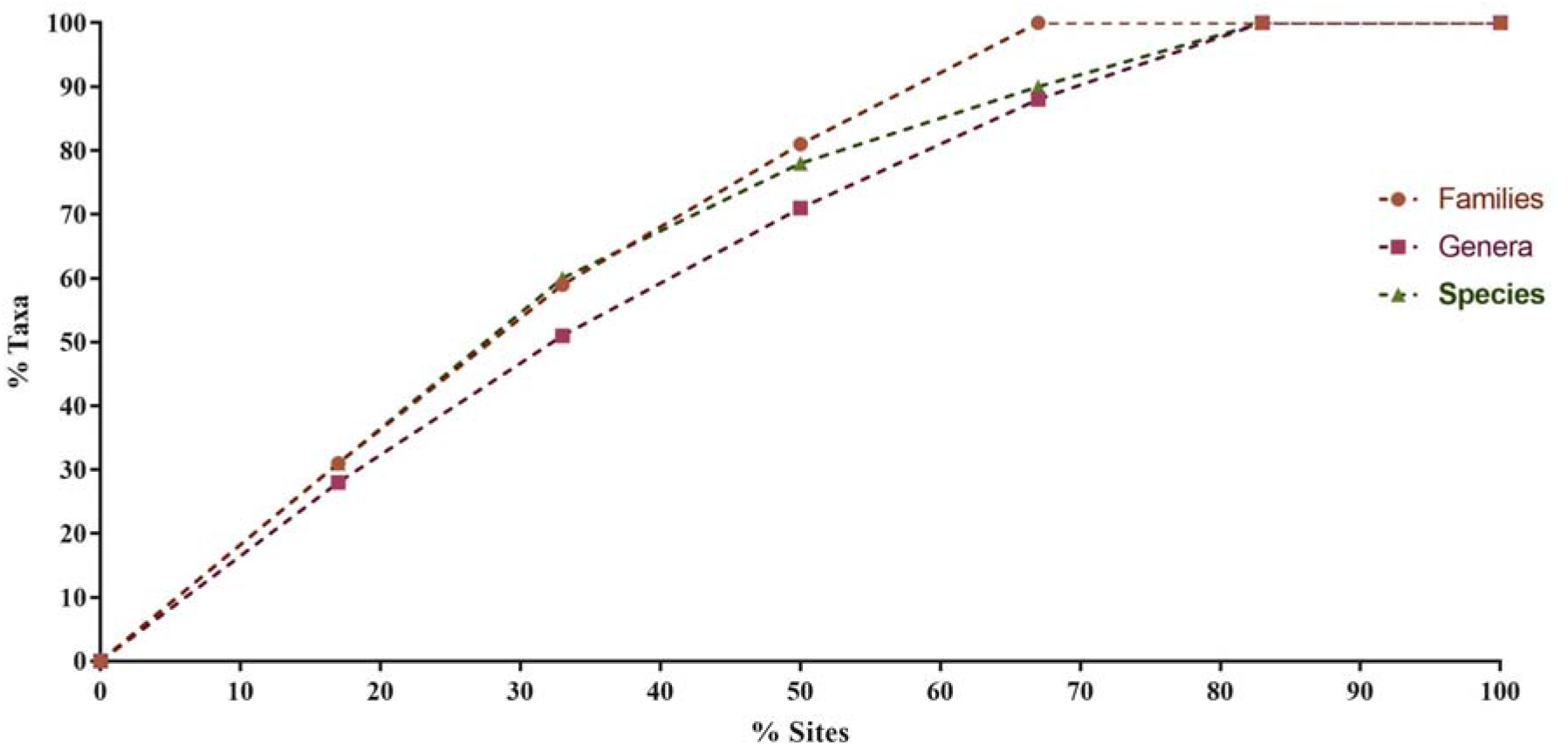
Accumulation curves of the number of taxa represented by the adding sites in a stepwise manner.

### Diversity indicator groups

After adjusting the linear regressions for the 11 families, the Lycosidae family was found to have the highest relationship with all species richness, with a regression coefficient of 79% (Table 4; Fig. 8 (a)). This family presented a significant relationship with the total number of species (n = 6, *P* = 0.0166). Subsequently, we added more families to the linear regressions to increase the value of the regression coefficient. We do this five times until a group of six families was reached (Table 4). The regression coefficient increased considerably after adding the first family; however, the scatter plot started at zero since Lycosidae and Araneidae were not present at all sites (Fig. 8 (b)). Adding the family Thomisidae to the regression analysis increased its regression coefficient considerably with a value of 97% (Fig. 8 (c)). The Lycosidae and Araneidae families are the richest in species. Although the Thomisidae family is not one of the richest, the three families together account for 45% of the total wealth. By continuing to add families to the analysis as expected, the regression coefficient increased to a regression coefficient of 99% (Table 4)

**Table 4.** Accumulation of families in the group of indicators, according to the family that increased the regression coefficient in each analysis.

| Family | Accumulated species | Regresion coefficient ($r^2$) |
| --- | --- | --- |
| Lycosidae | 8 | 0.797 |
| Araneidae | 19 | 0.943 |
| Thomisidae | 22 | 0.976 |
| Anyphaenidae | 24 | 0.982 |
| Dictynidae | 26 | 0.990 |
| Agelenidae | 29 | 0.998 |

**Figure 8.**
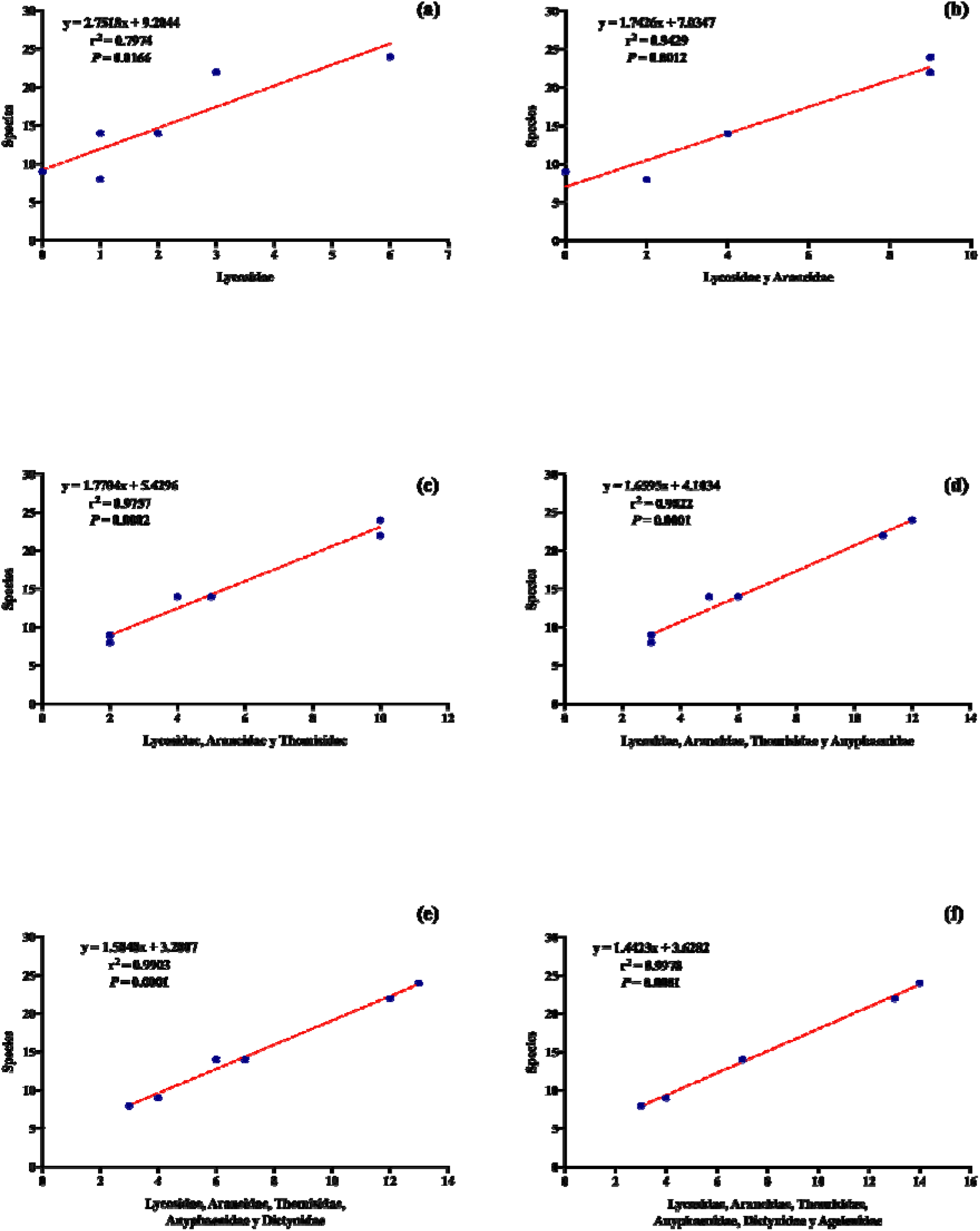
(a) Linear regression between Lycosidae and the total species richness of all localities, (b) Linear regression between an indicator group of two families and the total species richness of all localities. (c) Linear regression between an indicator group of three families and the total species richness of all localities. (d) Linear regression between an indicator group of four families and the total species richness of all localities. (e) Linear regression between an indicator group of five families and the total species richness of all localities. (f) Linear regression between an indicator group of six families and the total species richness of all localities

## Discussion

This work is the first arachnological study in a locality with geothermal influence in the north of Puebla and Mexico. A total of 371 organisms belonging to 49 species were collected. Although there are no data in the region that come from localities with geothermal characteristics, and there are no works evaluate spiders as surrogates, there are arachnological in other localities of the state of Puebla, such as Correa-Ramirez (2001) and Nieto-Castañeda (2000) where they recorder fewer species (29 and 31 respectively) than those in this study. Our results show a greater richness; however, we must consider that the environmental conditions were not the same. In the locality where we carried out our sample, there is a geothermal influence that could influence the spider’s composition.

In the evaluation of the integrity of our sample, the Chao 1 estimator indicated a missing in our sample of 47% of the richness, and the ACE estimator indicated a 30% missing. The integrity of our sampling was 53%, which is lower than that recorded in other studies (Cardoso, 2009; Cardoso et al., 2008; Maya-Morales et al., 2012; Sørensen, 2004; Sørensen, Coddington, & Scharff, 2002); this could be because our sampling effort was less or also because we only used one capture method, Cardoso (2009) suggests that a reasonable arthropod inventory reaches 50% of sampled species and an inventory full should reach between 70% and 80%; whereas, an exhaustive inventory reaches between 90% and 100%, but this would be impractical, since the sampling effort necessary to reach this level increases to a great extent.

The percentage of “singletons” obtained was 39%, a higher value than that found in other studies such as that of Cardoso (2008), who recorded 17% of “singletons” in an Oak forest in Portugal, Coddington (1996) recorded 29% in a forest in the United States, Toti (2000) recorded 34% in a forest in the United States and Sørensen (2004) who recorded 23% in a forest in Tanzania. The high percentage of “singletons” could be explained by subsampling or by various types of edge effect (Scharff, Coddington, Griswold, Hormiga, & Bjorn, 2003); As well as it is possible that some of these “singletons” are temporary, but this can only verify by sampling in all seasons (Toti et al., 2000), which is lacking in our study. Possibly, for this reason, our percentage of “singletons” was high.

There is a marked difference in the richness and abundance of spiders between localities. Although environmental variables were not measured, we can infer that these differences could be for several reasons, one of them and great importance is the felling of trees, Ozanne (2000) suggests that the felling of trees lowers the density of the vegetation, which causes a modification to the microclimates and due to this the richness and abundance of spiders are affected; In the sampled sites, we observe different levels of deforestation, some higher than others, another factor that affects the abundance and richness of spiders is the intensity of livestock foraging (Gibson, Hambier, & Brown, 1992), in all the localities we observe, foraging by cattle, sheep, and horses, that may be affecting the richness and abundance of spiders by modifying the density of vegetation and microclimates. These two factors must need considered in future work and measurements that allow us to evaluate spiders as possible indicators of the quality of management at these sites. Another aspect that has need considered is that in each locality, the influence of geothermal energy was different since some sampling points were further away than others from the nucleus of geothermal emanations, so this phenomenon could affect changes in abundance and spider diversity, as has already been reported in other studies in sites with similar conditions (Kovács et al. 2014; Jung et al. 2008, 2012).

The species accumulation curves did not reach the asymptote; this is similar to that reported in other studies that record the quality of spider samplings (Brennan, Majer, & Reygaert, 1999; Coddington, Young, & Coyle, 1996; RL Edwards, 1993; Samu & Lovei, 1995; Sørensen et al., 2002; Toti, Coyle, & Miller, 2000), to address this problem, is recommended to focus the studies on only some families or guilds of spiders and developing specific protocols for each family or spider guild (Sørensen et al., 2002); However, in our case, we not reach the asymptote this could also be due to the low number of samples obtained in our sampling.

Our study shows that spider genus richness data can infer spider species richness estimation. Similar results have been reported for other groups of terrestrial arthropods, such as the Diptera, Hymenoptera, and Coleoptera orders, which show that diversity substitutes can offer valuable information without having to perform identification to the species level (Báldi, 2003; Biaggini et al., 2007; Muñoz Gutiérrez, Rousseau, Andrade-Silva, & Charles Delabie, 2017); also, the use of diversity substitute taxa has been reported in the case of spiders (Cardoso et al., 2004; Lin et al., 2012).

Regression analyzes indicated that richness at the genera level has greater predictive power than families since it has a higher regression coefficient. This result is the same found by Cardoso et al., (2004) in Portugal, who found that the richness of genera has a greater predictive power compared to other taxonomic levels of spiders. That could be because a similar number of genera and species of spiders found at the sampling points worked and therefore have a higher correlation. However, genera as substitutes for diversity could cause problems, as it is sometimes difficult to even reach this level of identification. Cardoso et al. (2002), in a similar work in Portugal, also found that spider genera have greater predictive power than spider families; however, in contrast to this criterion, they analyzed the relationship of family groups with species richness, finding that family groups have a high predictive level of the species present in a given study site, unlike when using families separately.

Although higher taxa can help predict species richness, we need to consider some factors. Andersen (1995) and Cardoso et al. (2004) argue that the sampling effort and taxa’s geographic location can influence the predictions’ reliability. In contrast to this criterion, our sampling effort was much lower; however, the regression coefficient for the richness of genera in our analysis was similar to that found by Cardoso et al. (2004), suggesting that in our case, the sampling effort may not be a limiting factor; however, we need to do more tests in subsequent studies to assess whether sampling effort or vegetation types could influence the subrogation of higher taxa.

Higher taxa are not always enough to detect spider species richness changes (Cardoso et al., 2004). Another approach used to remedy it is the use of indicator taxa as substitutes. However, Cardoso et al. (2004b) suggest that more information could be lost using higher taxa subrogation. Groc et al. (2010) proposed a combined method, using higher taxa with indicator taxa at the species or morpho-species level. This method reduces the amount of taxonomic work linked to the identification of species and, at the same time, retains most of the biological information.

According to our study results, we can consider spider genera as suitable substitutes for choosing priority conservation sites, but not for spider families. This result is similar to that found by Cardoso et al. (2004). Cardoso (2004) suggests that precautions be taken when using this method. We need to take a conservative approach, trying to protect more sites than are expected to be necessary to represent all spider genera. That will ensure that the proposed sites protect a large proportion of species. However, using the genera as substitutes can also cause problems when making taxonomic identifications. Despite the difficulties that this method may have, we believe that it is a tool that will support rapid decision making in matters of the selection of priority sites for conservation.

Our work concludes that this tool can be very useful in a rapid study of biodiversity, especially when working with taxonomically complex organisms such as spiders. However, we consider that more complete studies should be carried out, evaluating the capture methods and vegetation types. However, this method can give us preliminary results for making decisions about conservation and environmental planning in geothermal sites, especially in those where the species present are unknown and when economic resources are limited. The spider species richness data is new for this area of Puebla in Mexico, and a question that arises in this research is, could spider richness be indicative of the geothermal condition of the area? Because the greatest richness of spider species was found in the closest areas with the geothermal influence that was the CCA and CUA sites. For this reason, we recommend future research to reinforce this research question.

## Conclusions

This study presents results showing that spiders can become surrogates of diversity in geothermal environments, allowing faster faunal studies on spiders and supporting management proposals for environments with these characteristics. We recommend more detailed studies considering vegetation and sampling efforts to confirm that this does not affect the predictions’ reliability.

We found a difference in abundance and diversity in each sampling site, which has a different geothermal influence, so we recommended carrying out studies to evaluate spiders as indicators of environmental contamination by heavy metals.

Within this study, we generate a fauna list of spiders for the geothermal area of Acoculco in Puebla, Mexico; however, the sampling integrity was low, so it is recommended in the future to carry out complete faunal monitoring of spiders to do so complement the list reported in this work.

## Acknowledgments

To the Autonomous University of Baja California for being a very important part of my academic and professional training. To the National Council of Science and Technology (CONACYT) for supporting me with the school scholarship (CVU 852359) that helped me carry out my studies. To the Crowdfunding platform “Experiment” for their trust and support to launch this project. To the Mexican Center for Innovation in Geothermal Energy (CeMIEGeo) for financing this research project.

